# Deconvolution and Phylogeny Inference of Diverse Variant Types Integrating Bulk DNA-seq with Single-cell RNA-seq

**DOI:** 10.1101/2025.01.24.634791

**Authors:** Nishat Anjum Bristy, Russell Schwartz

## Abstract

**Motivation:** Reconstructing clonal lineage trees (“tumor phylogenetics”) has become a core tool of cancer genomics. Earlier approaches based on bulk DNA sequencing (DNA-seq) have largely given way to single-cell DNA-seq (scDNA-seq), which offers far greater resolution for clonal substructure. Available data has lagged behind computational theory, though. While single-cell RNA-seq (scRNA-seq) has become widely available, scDNA-seq is still sufficiently costly and technically challenging to preclude routine use on large cohorts. This forces difficult tradeoffs between the limited genome coverage of scRNA-seq, limited availability of scDNA-seq, and limited clonal resolution of bulk DNA-seq. These limitations are especially problematic for studying structural variations and focal copy number variations that are crucial to cancer progression but difficult to observe in RNA-seq.

**Results:** We develop a method, TUSV-int, combining advantages of these various genomic technologies by integrating bulk DNA-seq and scRNA-seq data into a single deconvolution and phylogenetic inference computation while allowing for single nucleotide variant (SNV), copy number alteration (CNA) and structural variant (SV) data. We accomplish this by using integer linear programming (ILP) to deconvolve heterogeneous variant types and resolve them into a clonal lineage tree. We demonstrate improved deconvolution performance over comparative methods lacking scRNA-seq data or using more limited variant types. We further demonstrate the power of the method to better resolve clonal structure and mutational histories through application to a previously published DNA-seq/scRNA-seq breast cancer data set.

**Availability:** The source code for TUSV-int is available at https://github.com/CMUSchwartzLab/TUSV-INT.git

## Introduction

Cancer develops through an evolutionary process by which an initially healthy cell population successively accumulates mutations, generating genetic diversity that in turn creates opportunities for selection for more aggressive cell populations (clones) over the course of time [31]. While most such mutations are likely selectively neutral [36], chance deleterious mutations can lead cancers to select for phenotypic changes promoting tumor growth and progression [15]. Mutations causing pathway dysregulation might act through changes in gene copy number, molecular functions, and gene expressions, among other mechanisms. This resulting process of clonal evolution, together with the response it provokes in the tumor microenvironment, ultimately lead to the phenotype of tumor growth that defines cancer. Unraveling the enormous complexity of the process, though, has been the work of many years of effort into gathering diverse data sources on tumor genetics, function, and morphology and resolving their data into into coherent models of the process of tumor evolution and its relationship to disease progression [28].

Computational tools for reconstructing the process of clonal evolution in cancers from genomic data sources (“tumor phylogenetics”) have become a crucial part of cancer research, with their development emerging as its own vibrant subfield of research driven by and driving biotechnology development for profiling cancer progression [34] as well as helping to spur extensive study of somatic evolution beyond the cancer context [3]. The earliest methods for clonal lineage reconstruction relied on methods that predate the genomic era [25, 32]. Tumor phylogenetics only came into more widespread use as newer genomic technologies began to make cancer genetic variation data available at scale. The first widely used tools were designed to take advantage of bulk DNAseq (e.g., [4, 35]). The advent of scDNA-seq (scDNA-seq) for cancer studies [30] revolutionized clonal phylogenetics, with many methods today designed to us scDNA-seq data (e.g., [7, 33, 18]). However, scDNA-seq remains comparatively costly and technically challenging and so is still not widely used for large-scale cancer genomic studies or clinical practice [8]. Single-cell RNA-seq (scRNA-seq) is far more accessible and cost-effective [16], leading to efforts at scRNA-seq-based tumor phylogenetics [17, 20, 29], although such methods are challenged by the limited coverage of the genome they offer, poor resolution for fine-scale copy number changes, and limited ability to detect structural variations in particular.

Multiomic studies offer a potential path for leveraging advantages of distinct technology platforms. Multiomic methods have a long history in tumor phylogeny studies, including, for example, methods for combining bulk and scRNA-seq [22], bulk and scDNA-seq [26, 23], bulk DNA-seq and pregenomic fluorescent in situ hybridization (FISH) data [21], scDNA-seq and FISH [10], and scDNA-seq and scRNA-seq [6]. Particular value may be found in taking advantage of the comprehensive genomic coverage offered by bulk DNA-seq and the fine clonal resolution offered by scRNA-seq, both of which are now routinely available in low-cost commercial platforms, as recently adopted by some tumor phylogeny efforts [17, 40, 27]. Among these methods, PhylEx [17] and Canopy2 [40] use Bayesian methods to reconstruct tumor phylogenies from SNVs while Cardelino [27] is primarily an scRNA-seq method but can optionally use bulk sequence if provided a clonal guide tree. The benefits of this combination of technologies should be particularly pronounced as we move to develop more comprehensive models of how different variant types collectively contribute to tumor evolution, given the severe limitations of RNA-seq alone for profiling SNVs, CNAs, or SVs comprehensively. Recent work [11, 1] has shown the importance of looking at these classes of variants together, given their complementary value for accurately resolving clonal evolution and our growing appreciation for how they can each act in parallel in causing phenotypic changes that shape clonal evolution pathways [1, 39].

In the present work, we develop TUSV-int, a platform for clonal evolution studies integrating bulk DNA-seq and scRNA-seq to produce clonal evolution models that are unprecendented in their comprehensive accounting for fine clonal structure, genomic coverage, and diverse variant types (SNV, CNA, and SV). The work uses a general integer linear programming (ILP) framework of clonal lineage reconstruction in the presence of SVs [5] that we have previously adapted to provide similarly comprehensive variant coverage from bulk DNA-seq [11] and scDNA-seq [1] data. We validate TUSV-int in comparison to published alternative approaches, showing the value of considering multiple variant types and of combining bulk DNA-seq and sc-RNAseq in resolving clonal evolution accurately and comprehensively. We further demonstrate with application to a previously published DNA/RNA breast cancer data set [2] that TUSV-int is practical on real data and can yield improvements in clonal deconvolution and lineage reconstruction capable of driving novel biological discovery.

## Methods

We aim to reconstruct tumor clonal evolutionary histories, infer the SNV, SV, and CNA profiles of individual tumor clones, and determine their mixture fractions in samples of bulk DNA-seq using scRNA-seq data as a guide. We follow prior work of the original TUSV method [5] and variants [11, 1] in representing these variants for the formal problem statement. CNAs are represented as copy numbers for genomic regions, discretized into disjoint segments (bins). SVs are represented as pairs of genomic breakpoints that are adjacent in the cancer genome but non-adjacent in the reference genome. SNVs are represented by a single position in the genome. Allele fractions at SNV and SV positions are calculated as the ratio of reads supporting the alternate allele to those supporting the reference allele. We elaborate on these representations in defining problem variables in the following sections.

### Problem Statement

TUSV-int takes processed variant calls as input for both bulk DNA-seq (which may be WGS, WES, or targeted sequence data) and scRNA-seq. In bulk DNA-seq, we represent SVs with segmental mean copy numbers of paired breakpoint ends, CNAs with allele specific mean copy number for a set of discrete genomic segments, and SNVs with estimated mean copy numbers at the SNV positions. In scRNA-seq data, we represent CNAs with allele-specific mean copy numbers of genomic segments, which can be derived from existing copy number callers, such as Numbat [13] or SIGNALS [12]. Assume we are given: 1) *m* samples of bulk DNA-seq data covering *r* genomic segments and containing *l* SV breakpoints and *g* SNVs and 2) scRNA-seq data covering the same *r* genomic segments and corresponding to *n* tumor clones. Note that for computational efficiency, the variants used as input may be subsampled from the full variant set, a strategy for which we provide a more detailed quantitative justification in our prior work [1]. Given these inputs, we first construct an *m*×(*l*+*g*+2*r*) clonal variant matrix, *F*, from the bulk DNA-seq data. We then build an *n* × 2*r* clonal copy number matrix *C*^RNA^, from the scRNA-seq data. Additionally, we build an *l* × *r* breakpoint to segment mapping matrix, *Q*, identifying the segment in which each breakpoint end is located and an *l* × *l* breakpoint pair mapping matrix *G* identifying which breakpoints ends are paired to one another. The notations are described in Table 1 in more detail. Given the *F, C*^RNA^, *Q*, and *G* matrices, our goal is to estimate an *m* × *n* clonal mixture fraction matrix *U*, *n*×(*l*+*g* +2*r*) deconvolved clonal copy number matrix *C, n*×2*r* estimated RNA copy number matrix *C*^′^, and *n* × *n* single-cell RNA seq clone to bulk clone mapping matrix *M* . We do that by solving for a constrained optimization problem minimizing the following objective function:

**Table 1.** Key parameters and variables needed for TUSV_-INT_.

| Notation | Interpretation | Type |
| --- | --- | --- |
| $n$ | The number of clones | User defined |
| $m$ | The number of samples | Data derived |
| $l$ | The number of SV breakpoints | Data derived |
| $g$ | The number of SNVs | Data derived |
| $r$ | The number of DNA copy number segments | Data derived |
| $F \in \mathbb{R}_{\geq 0}^{m \times (l+g+2r)}$ | $F_{i,j}$: Average copy number of variant $j$ in sample $i$ | Constant |
| $Q \in \{0, 1\}^{(l+g) \times r}$ | $Q_{i,j} = 1$ iff breakpoint or SNV $i$ is in segment $j$ | Constant |
| $G \in \{0, 1\}^{l \times l}$ | $G_{i,j} = 1$ iff $i$ and $j$ are paired breakpoints, deriving from the same SV | Constant |
| $C^{\text{RNA}} \in \mathbb{Z}_{\geq 0}^{n \times (2r)}$ | $C_{i,j}^{\text{RNA}}$: Clonal copy number of CNA $j$ in clone $i$ | Constant |
| $U \in \mathbb{R}_{\geq 0}^{m \times n}$ | $U_{i,j}$: Fraction of clone $j$ in sample $i$ | Variable |
| $C \in \mathbb{Z}_{\geq 0}^{n \times (l+g+2r)}$ | $C_{i,j}$: Clonal copy number of variant $j$ in clone $i$ | Variable |
| $M \in \{0, 1\}^{n \times n}$ | $M_{i,j} = 1$ iff clone $i$ in the phylogeny represents RNA clone $j$ | Variable |
| $E \in \{0, 1\}^{n \times n}$ | $E_{i,j} = 1$ iff there is a directed edge from clone $i$ to clone $j$ | Variable |
| $A \in \{0, 1\}^{n \times n}$ | $A_{i,j} = 1$ iff clone $i$ is an ancestor of clone $j$ | Variable |
| $W \in \{0, 1\}^{n \times n \times (l+g)}$ | $W_{i,j,k} = 1$ iff breakpoint or SNV $k$ is mapped on the edge from clone $i$ to clone $j$ | Variable |

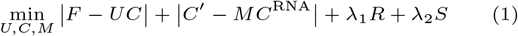

|*F* −*UC* | provides a measure of the quality of the deconvolution of the bulk data. |*C*^′^ − *MC*^RNA^ |provides a measure of consistency between the copy number estimates from deconvolved DNA-seq versus scRNA-seq. *R* is a regularization terms used to penalize for *L*1 cost of copy number distances along edges of a clonal phylogeny built from the inferred variants, used to bias the optimization towards a clonal deconvolution that yields a plausible evolutionary tree as assessed by an approximate minimum evolution cost. *S* is a measure of consistency between CNA and SV copy numbers. These terms and the constraint sets by which they are defined are described in more detail below.

### Algorithm

We formalize TUSV-int as a constrained optimization problem and solve for it with a coordinate descent algorithm, iterating between subproblems of the full optimization each of which we solve using integer linear programming (ILP). This is based on a strategy first developed for the original TUSV [5] and elaborated on in the more general TUSV-ext [11] for solving for tumor phylogenies integrating SNVs, CNAs and SVs from bulk-DNA sequencing data alone. The present work extends the TUSV-ext algorithm to accommodate the additional inputs, outputs, and constraints created by introducing scRNA-seq data.

We use coordinate descent to address the nonlinearity of our objective function in estimating the clonal mixture fraction matrix *U* and clonal copy number matrix *C*. The algorithm iteratively optimizes independently for *U* and *C*, starting with a random initialization of *U* . The variant copy number matrix *C* is estimated in two stages: 1) inference of the segmental CNAs from single-cell RNA sequencing data by estimating *M* and *C*^′^ and 2) estimation of the clonal SNVs and SVs constrained by the ILP formulation. We describe these in more detail in the following sections, focusing particularly on the extensions beyond TUSV-ext. We refer readers to the prior works [5, 11] for portions of the optimization that are largely unchanged, except where needed for clarity of exposition.

#### Estimating clonal mixture fraction matrix U

We define a set of ILP constraints to estimate *U* given *F* and *C*. The objective function is set to minimize the L1 norm of |*F* − *UC*|. Additionally, for each sample, we impose constraints so that the mixture fractions sum to 1 (Eqn. 2).

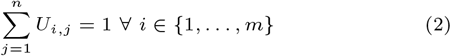

#### Estimating M and C^′^

Once *U* is estimated, we estimate the *n* × *n* bulk DNA-seq-to-scRNA-seq clone mapping matrix *M* and *n* × 2*r* estimated RNA copy number matrix *C*^′^ by minimizing the L1 norm of |*C*^′^ − *MC*^RNA^|. We impose constraints on *M* to ensure that each DNA clone corresponds to exactly one RNA clone (Eqn. 3, 4, 5).

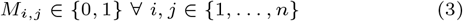

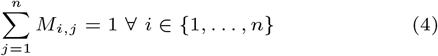

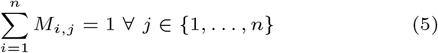

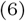

We then define the RNA copy number distance portion of the objective in Eqn. 1 using L1 distance as the following equations.

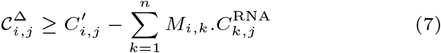

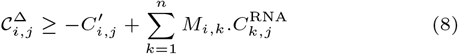

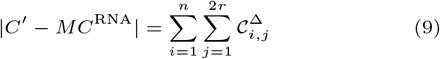

Simultaneously, we set the estimated *C*^′^ as the segmental CNA values of *C* (Eqn. 10).

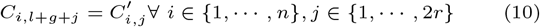

#### Edge and ancestry constraints

We model the underlying evolutionary history associated with the variants as a rooted directed binary phylogenetic tree T with *n* nodes corresponding to *n* clones, represented by a binary adjacency matrix *E*, where *e*_*i*,*j*_ = 1 if clone *i* is the parent of clone *j* for *i, j* ∈ {1, …, *n*}. To ensure that *E* represents a valid tree structure, we designate the first 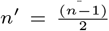 nodes to represent the leaves of the tree, *n*^′^ + 1 through *n* − 1 to represent the internal nodes, and node *n* to represent the root. *n*^′^ represents the number of internal nodes in T . We then impose the following constraints on the E matrix:

The root has no incoming edge and the leaves have no outgoing edge –

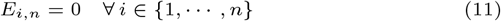

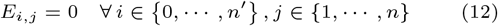

Additionally, following TUSV-ext [11], we define an ancestor matrix *A* ∈ {0, 1}^*n*×*n*^. *A*_*i*,*j*_ = 1 if clone i is an ancestor of clone *j*. We define the following two constraints to ensure that a node’s parent is its ancestor and a child node will carry all of its parent’s ancestors –

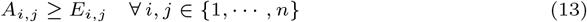

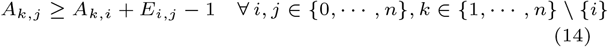

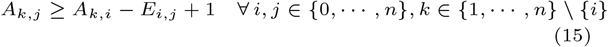

#### Copy number constraints

We define similar constraints to TUSV-ext [11] to ensure a tree structure on our estimated variant copy number matrix *C*. We assume that the root is a normal clone and therefore has no SNV/SV mutations, as well as normal diploid copy number at each genomic segment.

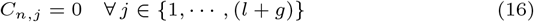

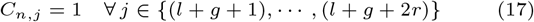

For algorithmic tractability, we constrain the maximum copy number to be a user-specifiable parameter *c*_max_.

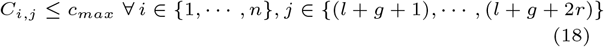

Additional constraints define the phylogenetic cost *R* as the sum of L1 distances of copy number differences across the edges of the tree. These are defined as in our prior work [11] and so are not reproduced here.

#### Dollo phylogeny on breakpoints and SNVs and breakpoint consistency

We apply constraints for the loss-supported Dollo phylogeny model, where a mutation can be gained only once, but lost multiple times. However, to favor consistency between CNA and SNV/SV data, we assume that SNV and SV losses are coupled with a loss of their corresponding genomic segments. Similarly, SNV and SV amplifications are constrained by their corresponding segmental amplifications, as described in TUSV-ext [11]. Additionally, we assign each breakpoint and SNV to an allele. This is achieved by introducing a binary matrix *D* ∈ 0, 1^(*l*+*g*)^, where *D*_*b*_ = 1 if and only if breakpoint or SNV *b* belongs to the first allele, and 0 otherwise.

Since each breakpoint belongs to exactly one segment, a breakpoint copy number should not exceed the copy number of its corresponding segment. For a breakpoint *b*, we impose the following constraints:

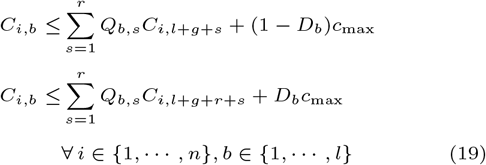

Finally, our second regularization term *S* in the objective function (Eqn. 1) enforces a relationship between the CNA segmental copy numbers and the breakpoint allele copy numbers. Since breakpoint copy numbers can be at most as large as the segmental copy numbers, the mixture copy number at the breakpoint position should also be at most the mixture segmental copy numbers. For a breakpoint *b* in sample *i*, we denote the ratio of the mixture copy numbers at the breakpoint or SNV position and the corresponding segment as *π*_*i*,*b*_.

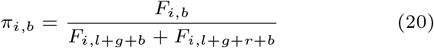

This ratio represents the sum of the amplification or deletion events of the mutation-containing allelle in comparison to the genomic segment as a whole. By minimizing the difference between the observed and estimated ratios, we align the estimated data with the observed mixture fractions. Following TUSV-ext [11], we include this minimization as a regularization term in the objective function to determine *S*.

### Choice of the regularization parameters

We set the regularization parameters to scale *R* and *S* so that they are comparable to the larger of the first two terms in the objective function. Specifically, we choose

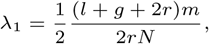

which is the ratio of the size of the first term to *R*, and

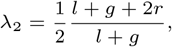

as in TUSV-ext [11], based on empirical observation that these formulas perform well in practice in the original TUSV and that solutions are minimally sensitive in the vicinity of these values.

### Availability

TUSV-int is implemented in Python 2 and utilizes the licensed Gurobi optimizer for ILP solution. The code is available at https://github.com/CMUSchwartzLab/TUSV-INT.git.

## Results

We first validated the accuracy and robustness of TUSV-int using simulated data and assessed its performance relative to TUSV-ext and Canopy2 [40]. While there are many tools available for tumor phylogenetics, there are no accepted standards for their specification and differences in inputs and outputs render direct head-to-head comparisons difficult. We compare to TUSV-ext to provide a direct assessment of the value of scRNA-seq in deconvolving bulk DNA-seq. A so-far unique feature of our TUSV-family methods is their simultaneous use of SNV, CNA, and SV data, and we therefore use Canopy2 to assess the comparative advantage TUSV-int gets from including additional variant types relative to a leading tool also using bulk DNA-seq and scRNA-seq.

Additionally, we evaluated TUSV-int’s effectiveness on real cancer patient samples. A full direct comparison of TUSV-int with other existing methods is again not feasible, as none of them utilize single-cell RNA sequences for clonal deconvolution from bulk DNA sequencing data using all of the variant types TUSV-int can access. We therefore again compare it with TUSV-ext, which reconstructs tumor phylogenies with bulk DNA sequences only with all of the same variant types, to assess what specifically we gain by adding scRNA-seq to the analysis.

Canopy2 [40] reconstructs tumor phylogenies from bulk DNA-seq and scRNA-seq based only on SNVs. While Canopy2 utilizes reference and alternate reads from bulk DNA-seq and scRNA-seq data, we use the copy numbers from scRNA-seq to guide our method. Canopy2 relies on assumptions of different mutation “activation” and “deactivation” rates in relating scRNA-seq data to clonal mixture fractions and mutation frequencies. In contrast, we do not explicitly model these rates. Instead, to ensure a fair comparison, we generate scRNA-seq reference and alternate read counts for Canopy2 assuming uniform activation and deactivation rates across all mutations and no sequencing errors. We achieve this by keeping the scRNA-seq reference and alternate read counts consistent with the clonal B-allele frequencies (BAFs) observed in the bulk mixture. For this purpose, we first extract clonal BAFs from the bulk mixture to represent the proportion of alternate allele reads for each clone. Next, for each single cell, we simulate reference and alternate read counts at each site by scaling the total read depth to 120x. These comparisons are meant only to assess the value of the unique combination of data sources (sequence types and variant types) used by TUSV-int and so we do not explore other scenarios where the variant types assumed by TUSV-int are not available.

### Evaluation with simulated data

To evaluate the performance of TUSV-int, we simulated bulk WGS data with SVs, SNVs, and CNAs based on known ground-truth tumor phylogenies, mutation rates, clonal populations, and their frequencies. Each simulation represented subclones from a single patient with two samples, using genomic segments from chromosomes 1 and 2. We began by assigning random clonal frequencies to the subclones. We then introduce genetic alterations, where we generate SNVs at uniformly random positions, and SVs with lengths drawn from a Poisson distribution with a mean of 5,745,000 bp, corresponding to the average SV length observed in the TCGA-BRCA cohort [5]. CNAs are generated using a relative probability distribution of 2:1:2:1 for amplifications, inversions, deletions, and translocations, respectively. To model sequencing errors, we generate segment-specific read counts using a Poisson distribution with a mean depth of 50. These read counts are used as the trial numbers for a binomial density that generates variant-associated read counts based on variant allele frequencies (VAFs). For each variant, we compute theoretical bulk DNA-seq copy numbers as a weighted sum across subclones based on their respective clonal frequencies. Using these copy numbers, we calculate variant allele frequencies (VAFs) for SNVs and breakpoints, as well as BAFs for CNAs. For transcriptomic simulations, we generate transcript counts using a Poisson density with a mean of 50 for at least 3 and at most 6 genes per genomic segment from the bulk DNA sequences for each clone. These counts are averaged and normalized for allele-specific single-cell RNA sequencing segment-wise CNA profiles.

We ran TUSV-int with the following parameter sets — (1) varying numbers of clones, *n* ∈ {3, 5, 7, 9} and (2) varying numbers of SNV mutations, *g* ∈ {25, 50, 100} — with each scenario having 3 iterations, 3 random restarts, 5000 seconds maximum compute time per iteration, 80 SVs, 40 SNVs (with the exception of 20 SNV for *g* = 25) and with the corresponding genomic segments for CNAs. Figure 1 shows the root mean square error (RMSE) of the true versus estimated clonal fraction matrices in these simulation settings.

**Fig. 1.**
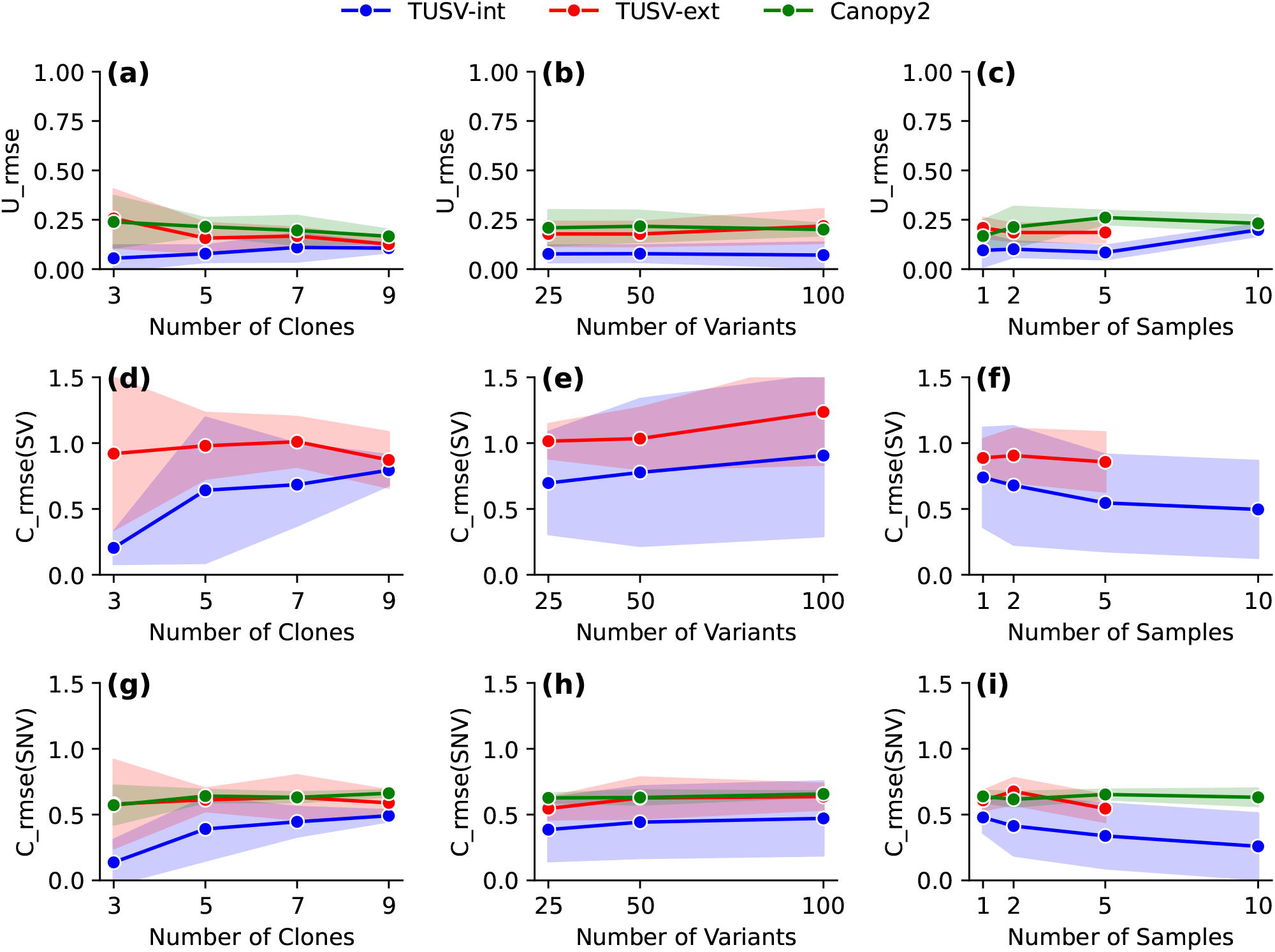
Results on simulation with different number of clones, samples and mutations. (a) - (c) shows mean squared error of the true vs estimated clonal mixture fractions (U) for TUSV-int, TUSV-ext and Canopy2. (d) - (f) shows the mean squared error of the estimated SVs for TUSV-int and TUSV-ext. (g) - (h) shows the mean squared error of the estimated SNVs for TUSV-int, TUSV-ext and Canopy2. We ran TUSV-int and TUSV-ext with m = 5000, t = 3, r = 3, sv ub = 80 and C = 120 (except for 25 SNVs, where C = 100). The solid lines represent the means of the simulation instances and shaded areas represent one standard deviation on either side of the mean. TUSV-intis evaluated at all data points. TUSV-ext and Canopy2 results were feasible for only for subsets of the data points, as described in the accompanying text. Note that clone 0 is assigned a small frequency in (c) despite not appearing in the TUSV-int phylogeny of (a) because it was inferred in one of the ten TUSV-int runs.

In all cases, TUSV-int outperformed our previous method, TUSV-ext, which relies solely on bulk DNA-seq. We observed a decrease in accuracy for TUSV-int as the number of clones increases (Figure 1 (a), (d), (e)), indicating that deconvolution becomes harder for a larger dataset. Inference of clonal SVs suffers more from increasing clone numbers than SNVs. Clonal frequencies show a smaller loss in quality with increasing numbers of clones than variant inferences. This is in contrast to TUSV-ext, which shows less sensitivity to clone numbers, although still worse performance than TUSV-int even under the most favorable conditions. This may correspond to the greater difficulty of accurately mapping scRNA-seq data to distinct DNA-seq clones as the clone number increases.

Increasing numbers of variants leads to less consistent trends between the two methods (Figure 1 (b), (e), (h)). Both methods show some loss of accuracy in SVs with increasing variant numbers, although more stable results for SNVs. Clonal frequencies are essentially stable for TUSV-int with increasing numbers of variants. TUSV-int remains consistently the better performer and the performance gap between the two methods is relatively stable across changes in variant numbers for the most part.

Increasing number of samples (Figure 1 (c), (f), (i)) generally led to improved solution quality by all measures, with the exception of the outlier of clonal frequency accuracy degrading for ten samples. We would generally expect more samples to lead to more effective deconvolution despite potentially increasing the size and difficulty of the optimization problem. There is a slight increase in separation between the two methods within increasing sample numbers, although not clearly distinguishable from chance. TUSV-ext did not yield results for ten samples within the allotted compute time. While improved runtime was not the goal of TUSV-int, the results may indicate an unanticipated value of scRNA-seq in more tightly constraining the problem and pruning the search space, thus accelerating ILP-type optimization.

We also evaluated Canopy2’s [40] performance on our simulated datasets, comparing to TUSV-int and TUSV-ext with respect to clonal frequency inference and SNV placement onto clones. For each simulation instance, we provided Canopy2 with the true number of clones and ran it using its default parameters. Fig. 1 shows Canopy2’s performance in green. Since Canopy2 does not make use of SVs, there are no Canopy2 results for Fig. 1(d,e,f). Root mean squared errors (RMSE) for clonal frequency inference and SNV placement with Canopy2 are generally comparable to those of TUSV-ext, with both notably worse than TUSV-int. The results confirm observations from prior TUSV variants that the addition of SV data leads to improved accuracy by other measures than just SVs, as SVs provide complementary information to other variant types that helps in correctly resolving the tree topology and inferring placement of all variants.

### Real breast cancer patient samples

To demonstrate TUSV-int’s performance on a true example of joint bulk DNA-seq and scRNA-seq data, we selected one dataset from an ER/HER2-positive breast cancer patient (BC03) among 11 patients studied by Chung et al. [2]. This dataset included three bulk WES samples collected from the primary tumor, metastatic lymph node, and blood. scRNA-seq was performed on 92 single cells from the primary tumor site and metastatic lymph nodes, and thus we included these two samples in our analyses while omitting the blood sequence for which no matched scRNA-seq is available. The bulk WES coverage was 100x for the tumor and metastatic lymph node, and 50x for the paired blood sample.

We preprocessed and called variants from the bulk WES data using the Nfcore/sarek pipeline [9, 14], aligning paired-end FASTQ files to the GRCh38 human reference genome. Raw sequencing reads were then trimmed and quality-controlled with fastp and FastQC, mapped using BWA-MEM2, deduplicated with GATK MarkDuplicates, and the base quality scores were recalibrated. Strelka2 [19], and CNVkit [38] were then used to call SNVs and allele-specific CNVs, respectively. We derived average allele-specific copy numbers from the total copy numbers reported by CNVkit, using the Strelka2 genotypes. We preprocessed the scRNA-seq data in a similar manner from the paired-end raw sequencing reads. Allele-specific CNA calls were then performed using Numbat [13]. Since Chung et al. [2] generated the RNA sequencing data using the SMARTer Ultra Low RNA Kit [2], we ran Numbat with the “SmartSeq” setting to accommodate the non-barcoded single-cell RNA sequences.

After obtaining SNVs and CNAs from bulk WES and allele-specific CNAs from scRNA-seq, we ran TUSV-int using 5 clones, 3 random restarts, 3 iterations, a time limit of 1500 seconds per iteration, 120 SNVs, and the corresponding average bulk CNAs alongside scRNA-seq-derived CNAs. For comparison, we also ran TUSV-ext using the same settings. We used the five scRNA-seq clones derived from Numbat, and calculated the average segmental copy number within each cluster to derive representative values as an input to TUSV-int.

Fig. 2 summarizes the results, comparing those between TUSV-int and the DNA-only TUSV-ext. TUSV-int inferred four clones (Fig. 2(a)) on 9 of its 10 runs, consistent with the results reported by Canopy2 for an optimal run on patient BC03’s data, and places the SNVs and SVs across all branches of the phylogenetic tree (Figure 2 (a)). On the other hand, TUSV-ext inferred five clones from the two samples (Fig. 2(b)), although its clone 0 and clone 1 have low support (Figure 2 (b)). The consistency between TUSV-int and Canopy2 results provides indirect support for the value of scRNA-seq to better resolve clonal structure. Clone 2 in both the trees have common mutations, however, they are placed differently in the two trees. We would expect the single-cell data available to TUSV-int to better identify which variants co-occur in the same clone and correct an error in placement in TUSV-ext. This leads to some notable changes in inference of mutation orders, for example in the observation that SNV mutations hitting the TP53 and Ras pathways (TP53BP1 and RASGRF2) and immune-related CNA mutation in TOLLIP occur at a late stage of progression following earlier mutations such as SNV mutation of SOX5 and CNA mutation of CD53. This is in contrast to the TUSV-ext result, which placed the mutations in clones 2 and 3 all as early mutations on two parallel clonal lineages. The distinction may be significant in interpreting the meaning of the results. For example, mutations affecting SOX5 and TP53BP1 can both promote proliferation in breast cancers through very different mechanisms [24, 37]. The TUSV-ext result would suggest these could be independently evolved ways of driving a similar phenotype in distinct clones while the improved TUSV-int result instead suggests they may be acting synergistically in the same cell lineage. Such a distinction can have important implications for identifying potential treatment options or predicting mechanisms of therapy resistance.

**Fig. 2.**
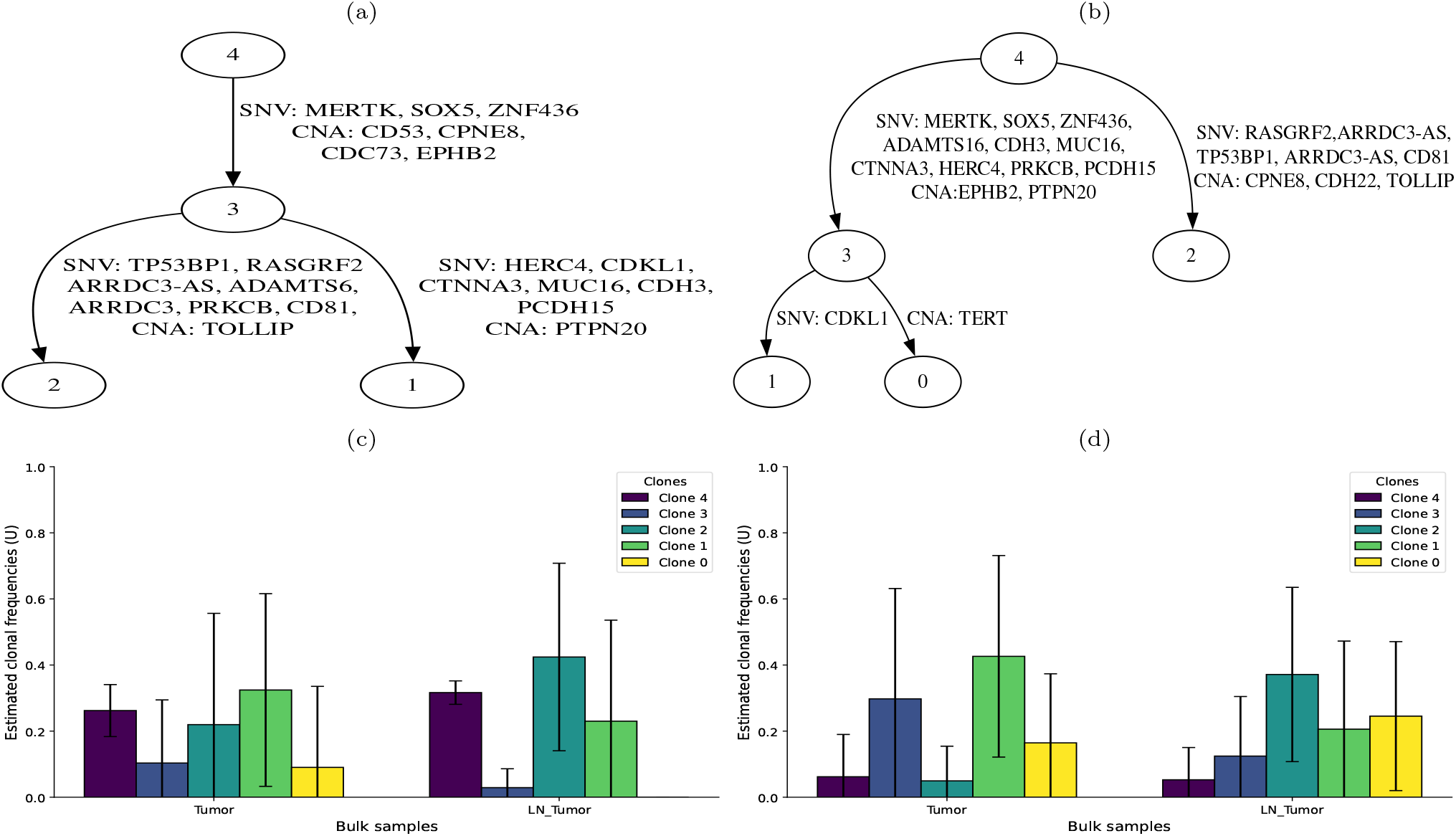
Results on ER/HER2 positive breast cancer patient BC03’s data. (a) Clonal tree inferred by TUSV-int with edges annotated by single-nucleotide variants (SNVs) and copy number alterations (CNAs). (b) Clonal tree inferred by TUSV-ext [11], also annotated with SNVs and CNAs. (c) Clonal mixture fractions in tumor and metastatic lymph node tumor (LN Tumor) samples inferred by TUSV-int. (d) Clonal mixture fractions in tumor and LN Tumor samples inferred by TUSV-ext. We ran TUSV-int and TUSV-ext with 120 subsampled SNVs, 3 iterations, 3 random restarts and 1500 second per iteration.

The two methods also yield fairly different inferences of clonal frequencies averaged across ten replicates (Fig. 2(c) and Fig. 2(d)), although the differences in the inferred clones themselves and their phylogenetic placements complicate interpretation. We would expect the scRNA-seq data to give TUSV-int a significant advantage in inferring accurate clonal frequencies, likely in addition to correcting errors due to misinferred clonal identities. TUSV-int and TUSV-ext both infer clone 1 to be the dominant clone in the primary tumor, although TUSV-int infers it to be substantially more diverged from the rarer clone 3 from which it descends. Both also infer clone 2 to be the dominant clone in the metastasis, making the revised placement of the clone potentially important in correctly characterizing the pathway to metastatic progression in this patient. Note that clone 0 is assigned a small frequency in the TUSV-int results despite not appearing in the TUSV-int phylogeny of Fig. 2(a) because it was inferred in one of the ten TUSV-int runs. There is little we can conclude about mechanism from a single patient, but the results illustrate how scRNA-seq data can lead to a more accurate model of the clonal evolution process and clonal dynamics across stages of progression, while taking account of how distinct types of genetic variants contribute to both phenomena.

## Discussion

We have developed a method, TUSV-int, for integrating bulk DNA-seq and scRNA-seq data for clonal phylogenies accommodating SNV, CNA, and SV variants. The work makes use of similar strategies for combining bulk DNA-seq and scRNA-seq to other recent works [17, 40] while handling a broader combination of variant types than any competing method. We accomplish this with a model built around an ILP framework for phylogenetic inference on heterogeneous data that offers comparatively easy extensibility to new data sources, variant types, and features of the biology. Validation on simulated data shows the advantages of combining both the two sequencing types and the three variant types in producing more accurate clonal deconvolution and lineage trees than do prior methods. Application to real data reinforces the importance of this combination in accurately resolving clonal evolution and revealing how different forms of variation can each contribute in parallel to the biology of cancer progression.

The work nonetheless leaves a number of avenues for improvement. Our method currently does not account for clones present in bulk DNA-seq but absent from single-cell RNA sequencing, a question we leave for future exploration. Improving computational efficiency is also a future concern, as ILP can be a computationally costly technique, in some cases necessitating subsampling of large data sets. ILP, despite its power for efficiently solving hard optimization problems, also has limitations compared to more common maximum likelihood and Bayesian approaches to phylogenetics. For example, combinatorial optimization methods like ILP provide less direct support for considering uncertainty in model inferences. It would have great value to the clonal lineage problems studied here and to and the field more generally to find new methods that can bring together the speed and versatility of ILP-like combinatorial optimization with the more principled handling of uncertainty of probabilistic models. There is also likely more advantage to be gained by considering ways to incorporate other data sources into the analysis, such as scDNA-seq, long-read sequencing, or data from blood-based sequencing (“liquid biopsy”). Finally, we have focused here primarily on methods development and the question is largely unexplored yet what the improved power of these methods can tell us about somatic variation and clonal evolution in cancer progression as well as in other diseases and normal aging.

## Author contributions statement

N.B. and R.S. conceived the method and study and wrote and reviewed the manuscript. N.B. implemented the code, executed the experiments, and analysed the results.

## Acknowledgments

We thank the Pittsburgh Supercomputing Center (PSC) and its ACCESS resource for providing compute resources used in the analyses described here. N.B. was supported in part by UPMC Enterprises through a Fellowship in Digital Health Innovation from Carnegie Mellon’s Center for Machine Learning and Health (CMLH). Research reported in this publication was supported by the National Human Genome Research Institute of the National Institutes of Health under award number R01HG010589. The content is solely the responsibility of the authors and does not necessarily represent the official views of the National Institutes of Health.

